# A high-throughput neutralization assay for yellow fever serodiagnostics

**DOI:** 10.1101/2021.12.16.472971

**Authors:** Madina Rasulova, Thomas Vercruysse, Jasmine Paulissen, Catherina Coun, Vanessa Suin, Leo Heyndrickx, Ji Ma, Katrien Geerts, Jolien Timmermans, Niraj Mishra, Li-Hsin Li, Dieudonné Buh Kum, Lotte Coelmont, Steven Van Gucht, Hadi Karimzadeh, Julia Thorn-Seshold, Simon Rothenfußer, Kevin K. Ariën, Johan Neyts, Kai Dallmeier, Hendrik Jan Thibaut

**Affiliations:** KU Leuven Department of Microbiology, Immunology and Transplantation, Rega Institute, Virology and Chemotherapy, Molecular Vaccinology & Vaccine Discovery, BE-3000 Leuven, Belgium; KU Leuven Department of Microbiology, Immunology and Transplantation, Rega Institute, Translational Platform Virology and Chemotherapy (TPVC), BE-3000 Leuven, Belgium; GVN, Global Virus Network; Virology Unit, Department of Biomedical Sciences, Institute of Tropical Medicine Antwerp, Antwerp, Belgium; Sciensano, Viral Diseases Service, Scientific Directorate of Infectious Diseases in humans, Brussels, Belgium; Division of Clinical Pharmacology, University Hospital, LMU Munich, Munich, Germany; Department of Biomedical Sciences, University of Antwerp, Antwerp, Belgium; Gene Therapy Division, Intas Pharmaceuticals Ltd., Biopharma Plant, Ahmedabad, Gujarat, India; Aligos Belgium, BE-3000 Leuven, Belgium; Unit Clinical Pharmacology (EKliP), Helmholtz Center for Environmental Health, Munich, Germany

**Keywords:** Yellow fever virus, serodiagnosis, reporter virus, PRNT, neutralization assay, high-throughput, Dengue virus, Zika virus, Japanese encephalitis virus

## Abstract

Quick and accurate detection of neutralizing antibodies (nAbs) against yellow fever is essential in serodiagnosis during outbreaks, for surveillance and to evaluate vaccine efficacy in population-wide studies. All this requires serological assays that can process a large number of samples in a highly standardized format. Albeit being laborious, time-consuming and limited in throughput, classical plaque reduction neutralization test (PRNT) is still considered gold standard for the detection and quantification of nAbs due to its sensitivity and specificity. Here we report the development of an alternative fluorescence-based serological assay (SNT^FLUO^) with an equally high sensitivity and specificity that is fit for high-throughput testing with the potential for automation. Finally, our novel SNT^FLUO^ was cross-validated in several reference laboratories and against international WHO standards showing its potential to be implemented in clinical use. SNT^FLUO^ assays with similar performance are available for the Japanese encephalitis, Zika and dengue viruses amenable for differential diagnostics.

**IMPORTANCE:** Fast and accurate detection of neutralizing antibodies (nAbs) against yellow fever virus (YFV) is key in yellow fever serodiagnosis, outbreak surveillance and monitoring of vaccine efficacy. Although classical PRNT still remains gold standard for measuring YFV nAbs, this methodology suffers from inherent limitations such as a low throughput and an overall high labor intensity. We present a novel fluorescence-based serum neutralization test (SNT^FLUO^) with equally high sensitivity and specificity that is fit for processing large number of samples in a highly standardized manner and has the potential to be implemented in clinical use. In addition, we present SNT^FLUO^ assays with similar performance for Japanese encephalitis, Zika and dengue viruses opening new avenues for differential diagnostics.

## Introduction

Yellow fever virus (YFV) is a mosquito-borne, positive-strand RNA virus that belongs to the genus *Flavivirus* within the family of the *Flaviviridae* and is the causative agent of yellow fever (YF). Other clinically important flaviviruses include dengue (DENV), Zika (ZIKV), West Nile (WNV) and Japanese encephalitis (JEV) virus (Gould & Solomon, 2008; Lindenbach, Thiel, & Rice, 2007; WHO, 2019). Despite the presence of a safe and very effective vaccine, that confers sustained immunity and lifelong protection after a single dose administration, YF still represents a major public health problem throughout the tropical areas of Africa, Central and Southern America (Barrett, 2017; Seligman, 2014; Theiler & Smith, 1937; WHO, 2019). In addition, insufficient vaccine coverage and international travel rise fear of YFV spreading to the Asian-Pacific regions where the competent mosquito vector is abundantly present and the human population is largely immunologically naïve to YFV (Butler, 2016; Kupferschmidt, 2016; Monath et al., 2016; Musso, Parola, & Raoult, 2018).

In the early and acute stages of disease, YF diagnosis is based on assessing patient’s clinical features in combination with conventional (end-point) or real-time reverse transcription polymerase chain reaction (RT-PCR) (PAHO, 2018; WHO, 2019), viral isolation or, in fatal conditions, immunohistochemical analysis to detect YF antigens in liver and other post-mortem tissues (De Brito et al., 1992; PAHO, 2018). In the later stages of infection, several serological methods are used to diagnose YF. Due to its simplicity to detect YFV-specific immunoglobulin M (IgM) and/or immunoglobulin G (IgG) antibodies, enzyme-linked immunosorbent assays (ELISA) have become the primary diagnostic tool worldwide (Adungo et al., 2016; Allwinn, Doerr, Emmerich, Schmitz, & Preiser, 2002; Basile et al., 2015; Domingo, Charrel, Schmidt-Chanasit, Zeller, & Reusken, 2018; PAHO, 2018). However, cross-reactivity with other flaviviruses or nonspecific reactivity often represents a major disadvantage in evaluating the infection in areas where other flaviviruses co-circulate (especially dengue and Zika viruses) (PAHO, 2018). Therefore, the more specific and sensitive plaque reduction neutralization test (PRNT), developed more than five decades ago (Spector & Tauraso, 1968), is currently still recommended as the “gold standard” assay worldwide. However, this assay suffers from several major disadvantages, including its duration, labor intensity, unsuitability for high-throughput settings, and the requirement of highly qualified and experienced staff to manually count plaque numbers (Simões et al., 2012).

Because of these technical drawbacks, there is an urgent need for a rapid, highly specific and robust surveillance and diagnostic tool. Such a method would help to recognize YFV outbreaks in an earlier stage, ease testing people with suspicion of YFV infection living in or returning from endemic regions, and hence to prevent viral spreading. In addition, in 2017 the WHO has launched a global strategy aiming to eliminate YF epidemics (EYE) by 2026 through vaccination of 1.4 billion people residing in YF-endemic areas and to contain outbreaks rapidly by the use of a fractional dose of YF17D (one fifth of the normal dose) (WHO, 2017). As it is still unclear how this fractional dosing will affect the life-long protection provided by YF17D (Barrett, 2020; Staples, Barrett, Wilder-Smith, & Hombach, 2020), large population-wide studies are required to monitor vaccine efficacy. Finally, during vaccine efficacy studies, it is also key to understand the impact of a new vaccine candidate on the effectiveness of existing vaccines, including YF17D (Alberer et al., 2015; Bovier, Althaus, Glueck, Chippaux, & Loutan, 1999; Clarke et al., 2016; Gil, González, Dal-Ré, & Calero, 1996; Haidara et al., 2018; López et al., 2016; Nascimento Silva et al., 2011; Sirima et al., 2021; Stefano et al., 1999; Stier, Weber, & Staples, 2012; Yvonnet et al., 1986). As such extensive studies require processing of a large number of serum samples, it is of essence to have a serological assay at hand that is robust and selective with improved turnaround and throughput properties.

Here, we describe an easy-to-use diagnostic method to quantify neutralizing antibodies using a fluorescently tagged YF17D as an alternative to classical PRNT. Our reporter-based neutralization assay is amenable for high-throughput screening with the possibility for automation, allowing detection of neutralizing antibodies in a large number of serum samples. This qualifies it as a powerful diagnostic tool in rapid identification of ongoing YFV outbreaks and assessment of vaccination efficiency in clinical trials, especially if only small serum volumes are available. Furthermore, quantification of neutralizing activity is fully automated, generating less subjective data than any manual counting method.

## Results

### Development of a high-throughput fluorescence-based seroneutralization test

To increase the speed of YF serodiagnosis it is key to implement a rapid and highly specific neutralization assay that is amenable for high-throughput screening and that is compatible with automated read-out and data analysis, ideally requiring only a limited volume of serum sample. Therefore, we developed an alternative, fluorescence-based seroneutralization test (SNT^FLUO^) to rapidly and reliably quantify YFV neutralizing antibodies in sera (Figure 1). To this end, we generated virus from a plasmid engineered to express YF17D together with mCherry (YFV/mCherry) as a translational fusion to the C protein (Figure 1a) (Kum et al., 2018; Mishra et al., 2020). The mCherry transgene remained stable up to at least six passages as demonstrated by RT-PCR fingerprinting performed on serially passaged YFV/mCherry in BHK-21J cells (Supplementary figure 1a,b).

**Figure 1.**
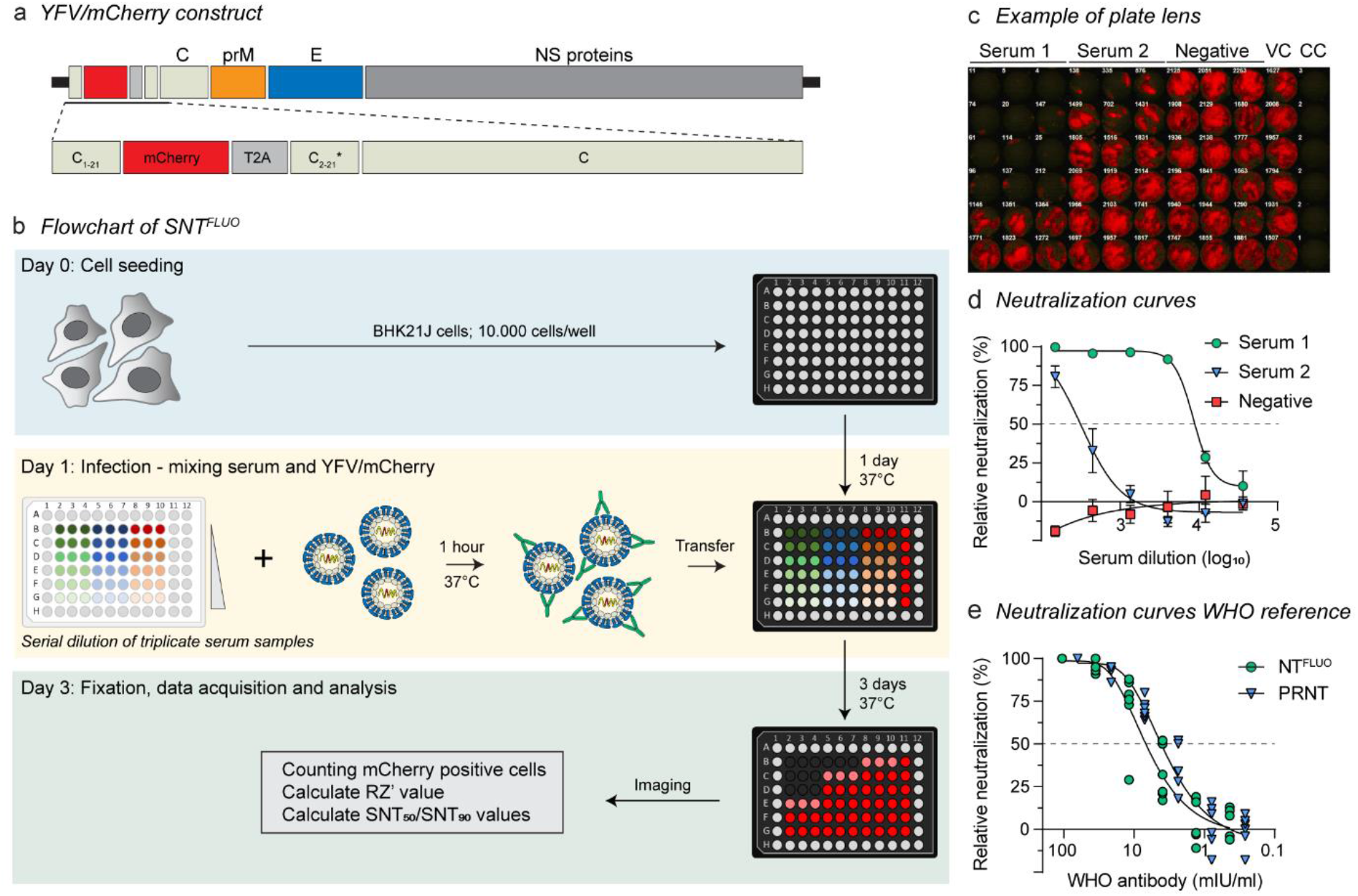
A high-throughput fluorescence-based seroneutralisation assay for YFV diagnostics. **a**, Schematic representation of YFV/mCherry. The reporter mCherry gene was inserted immediately downstream of codon 21 of YFV17D C gene and flanked by a *Thosea asigna* virus self-cleaving 2A peptide at the 3′ end, followed by a repeat of C gene codons with alternative sequence (C2-C21*). **b**, Assay flowchart of SNT^FLUO^. A detailed step-by-step bench protocol is provided as Extended Data File. YFV/mCherry was co-incubated for one hour with serially diluted sera in triplicate prior to infecting pre-seeded BHK-21J cells in a 96-well plate. Three days post-infection, cells were fixed, the fluorescent spots of infected cells were quantified and analyzed using a CTL ImmunoSpot® reader and Genedata screener software tool respectively. **c-d**, Representative image **(c)** and neutralization curves **(d)** of a counted plate containing 2 positive and 1 negative serum samples. **e**, Neutralization curves obtained by SNT^FLUO^ and PRNT on the WHO reference sample. Data are means ± standard deviations of three **(d)** or six **(e)** replicates.

As a first step towards assay set up, serially diluted YFV/mCherry was used to infect BHK-21J cells to identify the optimal balance between virus input (i.e. number of fluorescent cells, or spots), assay robustness (i.e. Z’ value) and assay endpoint (Supplementary figure 1c). On day 3, the highest Z’ value (i.e. >0.5) was obtained at the lowest virus dilution, immediately below a saturation point (i.e. ∼2000 spots/well of a microtiter plate). Latter parameters were chosen for further assay development and validation.

A flowchart for our SNT^FLUO^ in a 96-well format is depicted in Figure 1b. A detailed step-by-step bench protocol is provided as Extended Data File. Briefly, sera were serially 1:3 diluted in triplicate in round-bottom 96-well plates and co-incubated with YFV/mCherry for 1 hour at 37°C. Subsequently, the virus/serum mixtures were added to pre-seeded BHK-21J cells and further incubated for another three days. After fixation, the number of spots were quantified using a CTL ImmunoSpot® Series 6 Ultimate reader (Figure 1c) and data analyzed using Genedata Screener software package. In this way, assay statistics and dose-response curves are automatically calculated to determine the serum dilution fold that reduces 50% of the number of spots (SNT_50_) (Figure 1d).

For further initial validation and benchmarking of our reporter assay, we performed both SNT^FLUO^ and conventional PRNT on the WHO International reference sample (standardized monkey serum preparation containing YFV nAb) (National & Control, 2011) for a direct head-to-head comparison of the respective neutralization results (Figure 1e). EC_50_ values obtained by either assay were very much comparable and in the same range: 6.4 mIU/ml (95% confidence interval 5.2 to 7.8) and 4.0 mIU/ml (95% confidence interval 3.4 to 4.7) for the SNT^FLUO^ and PRNT respectively, as first evidence for similar performance of SNT^FLUO^ and PRNT. In conclusion, our results show that the YFV/mCherry reporter assay could serve as an equally sensitive, easy-to-use alternative for the gold standard PRNT.

### Assay validation by benchmarking against other neutralization tests

To further validate our SNT^FLUO^, we quantified neutralizing antibody levels in a large set of historical serum samples from previous vaccination studies with YF17D in mice, hamsters and non-human primates (NHP). For this purpose, we used several methods, including the here described SNT^FLUO^, conventional PRNT and another 96-well based neutralization assay that uses CPE as read-out (SNT^CPE^) (Mishra et al., 2020; Müller et al., 2017). PRNT and SNT^FLUO^ were performed on 67 serum samples (Figure 2a) and SNT^CPE^ and SNT^FLUO^ on 113 serum samples (Figure 2b), including only samples with nAb titers above LLOQ in both assays for calculation of correlation coefficients in regression analysis. The strongest correlation was observed between PRNT_50_ and SNT_50_^FLUO^ with R^2^ = 0.68 (Figure 2a, left panel). A weaker correlation was observed between SNT_50_^FLUO^ and SNT_50_^CPE^ with R^2^ = 0.57 (Figure 2b, left panel). Bland-Altman analysis was used to estimate the degree of agreement between assays, and to reveal any possible bias between their mean differences (Giavarina, 2015). In line with our regression analysis, the smallest bias and 95% confidence intervals (i.e. limits of agreement, LOA) was found between PRNT_50_ and SNT_50_^FLUO^ values with an average bias of 0.19 ± 0.42 (LOA from -0.63 to 1.0) (Figure 2a, right panel); whereas a markedly increased bias of -0.54 ± 0.43 (LOA from -1.4 to 0.31) was observed in the comparison of the SNT_50_^CPE^ and SNT_50_^FLUO^ values (Figure 2b, right panel). These results further confirm that SNT^FLUO^ yields neutralization results that are very comparable to those obtained by PRNT.

**Figure 2.**
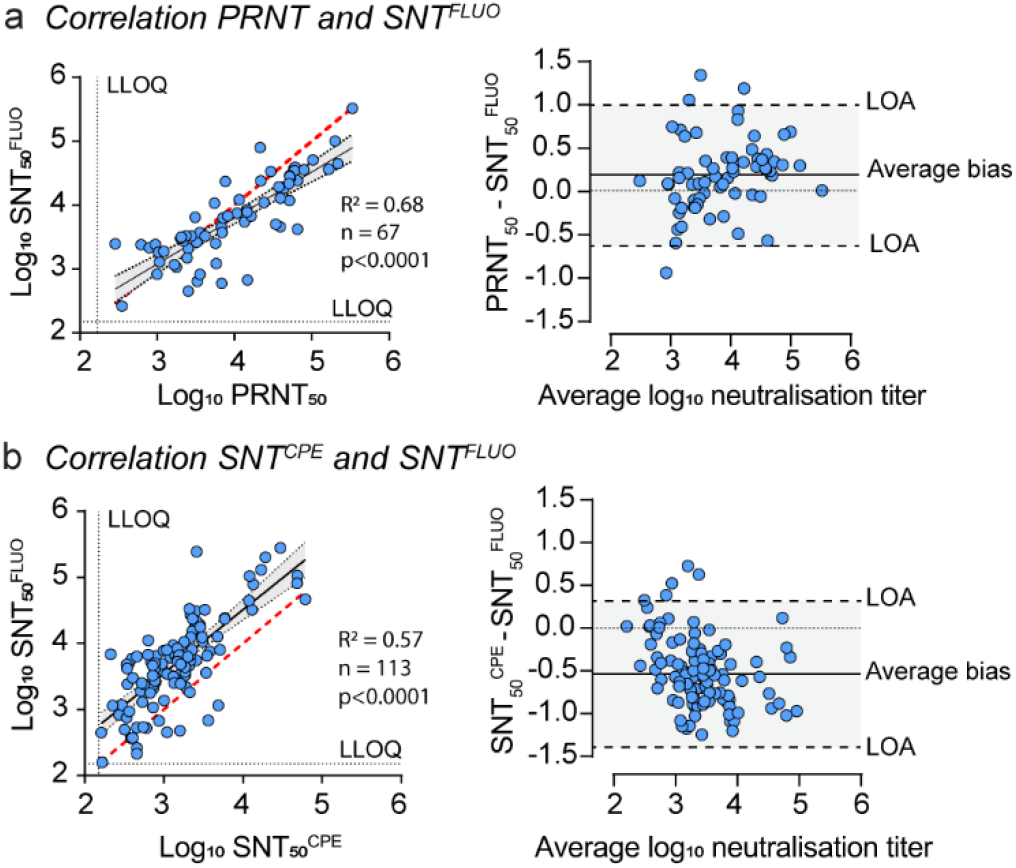
Benchmarking against other seroneutralisation assays. **a-b**, Correlation analysis of PRNT and SNT^FLUO^ **(a)**, and SNT^CPE^ and SNT^FLUO^ **(b)**. Left panels, linear regression analysis to calculate correlation coefficients. The Pearson correlation efficiency R^2^, number of tested sera (n) and *p*-values are indicated. Perfect correlation is indicated by the red dashed line whereas correlation between SNT_50_^FLUO^ and PRNT_50_ or SNT50FLUO and SNT50CPE is indicated by a black solid line. 95% confidence intervals are indicated by grey shaded areas. LLOQ - Lower limits of quantification. Right panels, Bland-Altman analysis to estimate the degree of agreement between assays. Differences between PRNT50 and SNT_50_^FLUO^ and SNT_50_^CPE^ and SNT^50FLUO^ values are compared with their average log_10_ neutralisation titer. The lines of no bias (dotted line), average bias (solid line), and 95% confidence intervals (i.e. lower and upper limits of agreements (LOA, dashed lines)) are shown. Data are means from three replicates.

### Assay cross validation by reference laboratories

A set of serum samples was prepared for cross-validation by the National Reference Center for Arboviruses at the Institute of Tropical Medicine in Antwerp (PRNT, ISO 15189) and Sciensano, Viral Diseases Service in Brussels (rapid fluorescent focus inhibition test (RFFIT), ISO 17025), two accredited reference laboratories in Belgium. To this end, serum samples from non-vaccinated (n=4) and YF17D-vaccinated (n=6) NHP were serially 1:10 diluted (1:10^1^ – 1:10^4^) and assessed blindly for the presence of nAbs in the three different laboratories using their respective assays (Supplementary table 1 and Figure 3). Six technical repeats of in total 34 serum samples were used to obtain dose response curves from which 50% neutralizing activities (EC_50_) were calculated. For correlation analysis between the three different assays, all nAb titers within the respective lower and upper quantification limits of each sample were included (Table S1). The strongest correlation was observed between SNT_50_ ^FLUO^ and PRNT_50_ values (Figure 3) with R² = 0.85, whereas R² = 0.78 was observed between SNT_50_ ^FLUO^ and RFFIT_50_. These results indicate that the SNT_50_^FLUO^ has the potential to be implemented in reference laboratories, including for high-throughput serodiagnostics, as a highly sensitive alternative for conventional time-consuming assays.

**Figure 3.**
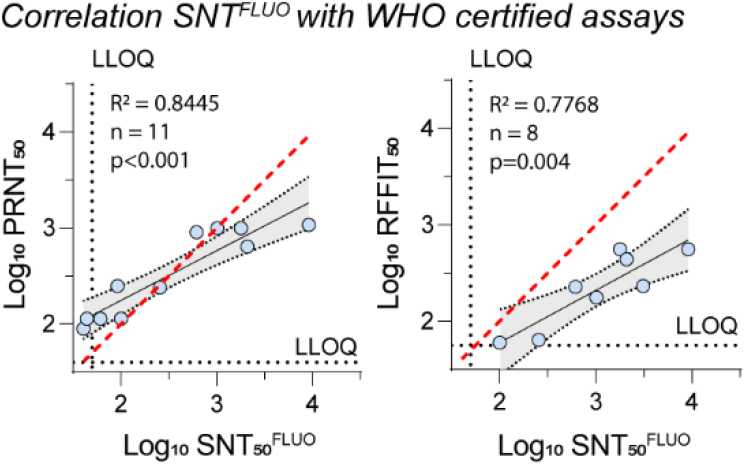
SNT^FLUO^ cross validation by reference laboratories. YF17D-vaccinated (n=6) and unvaccinated (n=4) monkey sera were serially 1:10 diluted (1:10^1^ – 1:10^4^) and assessed for the presence of neutralizing antibodies by the three different laboratories using their respective assays. The EC_50_ values for each serum sample between the upper and lower quantification limits was used for correlation analysis (Table S1). Data were analysed by linear regression to calculate correlation coefficients. The Pearson correlation efficiency R^2^, number of tested sera (n) and *p*-values are indicated. Perfect correlation is indicated by a red dashed line whereas correlation between the different assays is indicated by a black solid line. 95% confidence intervals are indicated by grey shaded areas. LLOQ - Lower limits of quantification. Data are the means of six (ITM and our laboratory) or two (Sciensano) replicates.

### Assay selectivity

Assay selectivity of SNT^FLUO^ was assessed using four groups of potentially cross-reactive sera from previous studies in which mice were vaccinated for either YF17D (n=8), Japanese encephalitis virus (JEV, n=8), Zika virus (ZIKV, n=9) or dengue type 2 virus (DENV2, n=5) (Figure 4a). All 30 serum samples were assessed for selectivity and cross-reactivity using similar fluorescence-based SNT^FLUO^ assays using four distinct reporter viruses, YFV/mCherry, DENV2/mCherry, ZIKV/mCherry and JEV/GFP (Figure 4b). DENV2/mCherry (Li et al., 2020) has been described earlier; ZIKV/mCherry and JEV/GFP were generated accordingly for this study. All test sera specifically neutralized only their corresponding cognate reporter virus without detectable cross-neutralizing activity towards the other flavivirus reporters (Figure 4c and supplementary figure 2). These data indicate that the YFV/mCherry SNT^FLUO^ is highly specific without notable flavivirus cross-reactivity. Furthermore, a panel of similar fluorescence-based neutralization tests may be added to the current diagnostic repertoire to aid rapid serological identification and differentiation of flavivirus infections.

**Figure 4.**
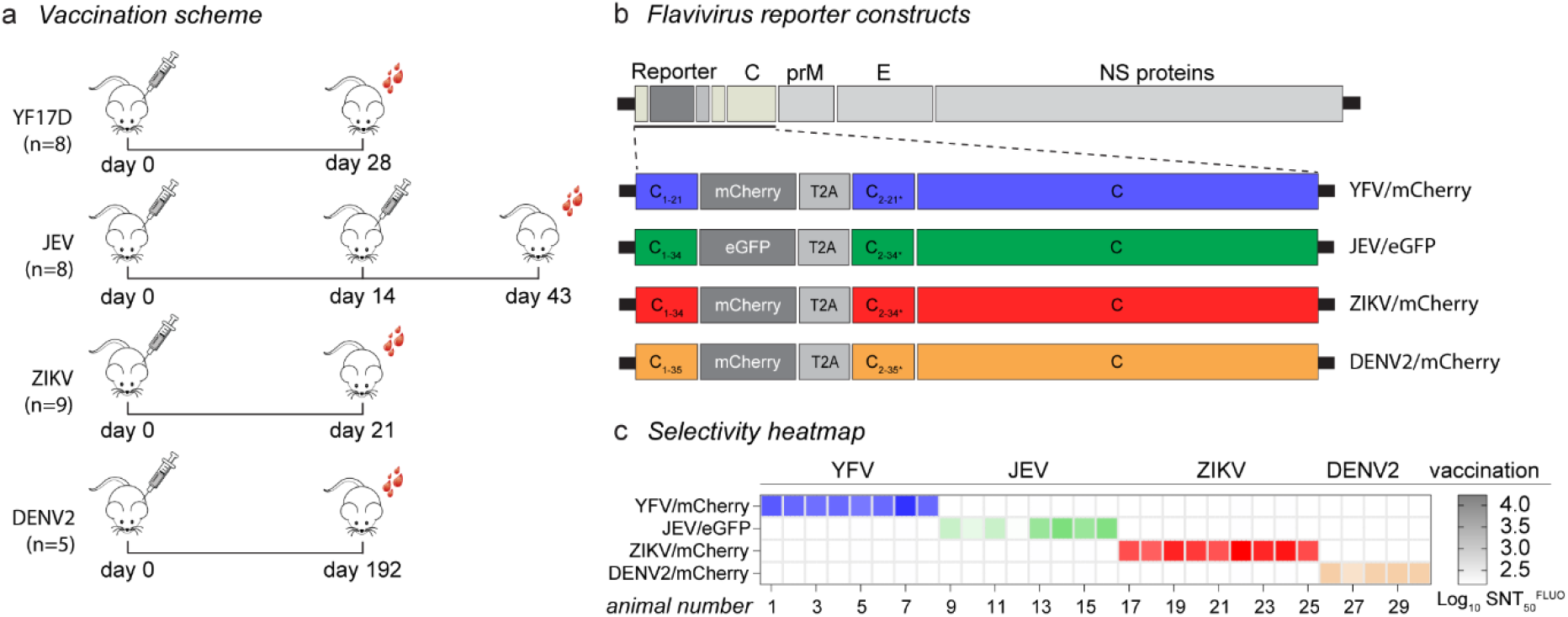
Selectivity of SNT^FLUO^ against other flaviviruses. **a**, Schematic representation of mice vaccination scheme for YF17D (n=8), JEV (n=8), ZIKV (n=9) and DENV2 (n=5), and day of serum collection (total of 30 serum samples). **b**, Schematic representation of YFV/mCherry, JEV/eGFP, ZIKV/mCherry and DENV2/mCherry. Reporter viruses were generated in a similar way as for YFV/mCherry, with the exception of extended and alternative codons of the C gene respectively flanking both ends of the fluorescent tag sequence (34 for JEV and ZIKV, and 35 for DENV2). **c**, Selectivity heatmap of neutralizing titers (SNT_50_^FLUO^) of the 30 serum samples tested using the 4 reporter flaviviruses. Data are the means of three replicates. Heatmap gradient indicates log_10_ NT_50_^FLUO^ values.

### Implementation of SNT^FLUO^ for high-throughput screening (HTS)

To explore whether our novel SNT^FLUO^ approach is fit for high-throughput serodiagnosis, we tested approximately 2000 serum samples from different organisms (mouse, hamster, pig, monkey, and human), either unvaccinated or vaccinated with YF17D. In total more than 600 plates were analyzed for robustness by calculating Z’ values, coefficients of variation (CV%) and signal-to-background ratios (S/B) (Figure 5a). The Z’ value achieved for the majority (>90%) of the plates was > 0.5, with a variability of <20% and S/B between 200 and 3500. About 1% of plates with a Z’ <0.1 needed to be rejected, whereas plates with a Z’ value between 0.1 and 0.5 (7%) were subjected to visual inspection (by fluorescence microscopy) to identify outliers, or possibly, to be rejected (Figure 5a, left panel, orange and red circles). Notably, poor assay robustness corresponded with a decreased in virus titer (reduced spot counts in untreated virus control wells), as expected for YF17D stocks during long-term storage at -80°C (Supplementary figure 1c), requiring back titration and adjustment of the infectious input material as performed during initial assay set-up. Finally, suitability of SNT^FLUO^ to monitor YF17D vaccination efficiency in larger population-wide studies was assessed. Therefore, we quantified the levels of nAb in matched serum samples (total of 727 sera) from a large cohort of subjects prior and after YF17D vaccination (at 2 different time points) (Figure 5b). Additionally, these 249 prior vaccination samples were used to accurately determine the limit of detection of our assay (i.e. 1:10). Altogether, the SNT^FLUO^ assay was capable of reliably and rapidly diagnosing the absence, or likewise seroconversion to YFV-specific nAbs, further demonstrating its potential to be used in a clinical setting where a fast sample turnaround is highly desirable.

**Figure 5.**
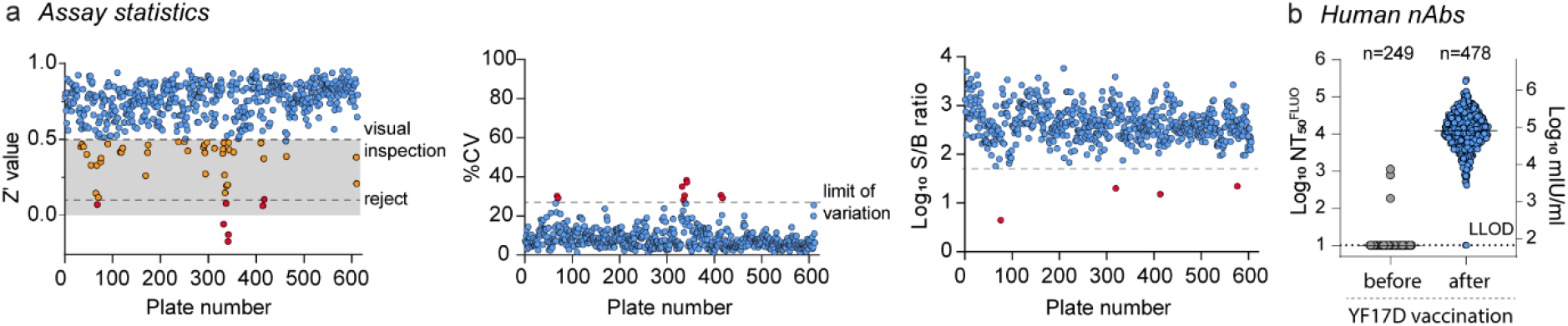
High-throughput performance of SNT^FLUO^. **a**, 600 assay plates containing approximately 2000 serum samples were analyzed for assay robustness by calculating Z’ values (left panel), coefficients of variation (CV%, middle panel) and signal-to-background ratios (S/B, right panel). Grey area in the left panel indicates Z’ score between 0 and 0.5. Plates with a Z’ score between 0.5 and 0.1 are subjected to visual inspection by fluorescence microscopy (orange circles). Plates with a Z’ < 0.1 are rejected (red circles). **b**, YF17D vaccination efficiency in a larger population-wide study prior and post vaccination. Blood was collected on day 14 and day 28 post vaccination and assessed for their neutralizing activity. Data are presented as Log_10_ NT_50_^FLUO^ (left Y-axis) or transformed to Log_10_ mIU/ml (right Y-axis). Data are the means of three replicates. LLOD - Lower limits of detection.

## Discussion

Fast and accurate quantification of YFV-specific nAbs plays a key role in YFV serodiagnosis, surveillance of large cohorts or population-wide monitoring of vaccine efficacy. Currently, the highly sensitive and selective PRNT assay is considered gold standard for evaluating the presence of YFV-specific nAbs, and superior to any other serological method established (Niedrig, Lademann, Emmerich, & Lafrenz, 1999). PRNT assays, however, suffer from several drawbacks including its labor intensity, duration and low throughput. Furthermore, PRNT assay robustness is difficult to control and nAb titers thus obtained may vary widely among laboratories. In this study, we developed a fluorescence-based yellow fever virus neutralization (SNT^FLUO^) assay as a high-throughput, rapid and easy-to-use alternative to the traditional PRNT.

We characterized the sensitivity and selectivity of the YFV/mCherry reporter SNT^FLUO^ by using a WHO reference standard and more than 2000 historical serum samples originating from several species backgrounds, including mice, hamsters, pigs, NHP and humans. A head-to-head comparison of SNT_50_ and PRNT_50_ values obtained by SNT^FLUO^ and PRNT, respectively, further demonstrated a similar sensitivity and good sample to sample correlation between both assays. Cross-validation of a set of NHP serum samples by two accredited reference laboratories in Antwerp (ITM) and Brussels (Sciensano) further underlined the possibility of adding the SNT^FLUO^ to the current YF serodiagnosis assay repertoire.

Due to its complexity and requirement of highly trained personnel to warrant reproducibility of data, PRNT assays are currently confined solely to accredited reference laboratories (Domingo et al., 2018). To overcome this bottleneck, there is a big need for an assay amenable to automation, including a fully automated data analysis pipeline. To this end, we used a fluorescent reporter yellow fever virus expressing an mCherry reporter which was chosen because of its small monomeric size, excellent brightness and superior photostability (Shaner, Steinbach, & Tsien, 2005). These properties render the SNT^FLUO^ assay highly suitable for fluorescence-based quantification by automated imagers. In addition, by using predetermined scanning and counting settings, user-dependent biases are kept to a minimum. In parallel, assay performance statistics are automatically calculated allowing the accurate monitoring of assay quality.

Compared with the PRNT assay, our SNT^FLUO^ has shortened the assay duration to four days, decreased the volume of serum that is required, facilitated assay manipulation (e.g. by omitting overlays) and increased the testing capacity to high-throughput. Although this assay has been developed for 96-well plate format, upscaling to a 384-well plate format should be possible further increasing its throughput. In addition to its high sensitivity, we also showed that the YFV/mCherry SNT^FLUO^ assay exhibits exceptional selectivity with no detectable cross-neutralization by potentially cross-reactive sera from animals vaccinated for either ZIKV, DENV2 or JEV. However, the level of possible cross-reactivity and cross-neutralization still needs to be explored in a clinical setting where multiple heterotypic infections can occur. In addition, we report on several other flavivirus reporter-based SNT^FLUO^ assays, including for ZIKV, DENV2 and JEV. Similar to YFV/mCherry no cross-reactivity was observed between the different sera. Although these assays still need to be further characterized and validated, our data already suggest that they could be implemented in a similar way as the YFV/mCherry SNT^FLUO^ assay. The availability of such assays with no cross-reactivity could be of great benefit for differential serology to accurately and rapidly identify a specific flavivirus infection which is especially challenging in regions where several flaviviruses co-circulate. Finally, similar reporter assays could be optimized towards high-throughput antiviral screening campaigns, either as single (Li et al., 2020) (Zhang et al., 2020) or, by choice of compatible fluoroprotein reporters, multiplexed antiviral screens possibly allowing to identify pan-flavivirus antivirals.

In conclusion, our SNT^FLUO^ assay offers a powerful tool to rapidly and selectively identify YFV-specifc nAb. Furthermore, its sensitivity, robustness and suitability for high-throughput screening makes it a valuable alternative to conventional PRNT.

## Materials and methods

### Cells

Baby hamster kidney fibroblasts (BHK-21J) and African green monkey kidney (Vero E6) cells were maintained in Minimum Essential Medium (Gibco) supplemented with 10% fetal bovine serum (HyClone), 2 mM L-glutamine (Gibco), 1% sodium bicarbonate (Gibco), 1x MEM non-essential amino acids solution (Gibco), 10 mM HEPES (Gibco) and 100 U/ml penicillin-streptomycin (Gibco) (hereafter referred as seeding medium) and incubated at 37°C. All assays were performed in the same medium but containing 2% fetal bovine serum (assay medium).

### Plasmid construction

Plasmid construction of the yellow fever 17D vaccine strain (pShuttle-YFV/mCherry) and Dengue virus type 2 NGC strain (pShuttle-DENV2/mCherry) reporter viruses stably expressing the red fluorescent protein mCherry have been reported previously (Li et al., 2020; Mishra et al., 2020). In a similar manner, Zika (pShuttle-ZIKV/mCherry) and Japanese encephalitis (pShuttle-JEV/eGFP) reporter viruses were generated for the current study (schematic representation in Figure 1 and Figure 4). In brief, cDNA of ZIKV strain BeH819015 (Mutso et al., 2017) was first inserted in pShuttle. This construct together with the previously described pShuttle plasmid containing JEV strain SA14-14-2 (Mishra et al., 2020) were next used to generate ZIKV and JEV reporter viruses expressing mCherry or eGFP respectively, as a translational fusion to the N-terminus of the C protein. Using standard molecular biology techniques and homologous recombination in yeast (strain YPH500), the synthetic DNA fragments encoding codons 2 to 236 of mCherry (GenBank accession no. AY678264.1) or codons 1 to 238 of eGFP (GenBank accession no. HI137399.1) were inserted immediately downstream of codon 34 of the C gene of the corresponding virus. The reporter gene is flanked by a BamHI restriction site at its 5′ terminus and by the ribosome-skipping 2A sequence of *Thosea asigna* virus (Demidenko, Blattman, Blattman, Greenberg, & Nibert, 2013) at the 3′ end. C gene codons 2-34* were repeated with an alternative codon usage to avoid recombination during virus replication. The plasmids were recovered from yeast, transformed into Epi300T (Epicenter) competent *E. coli* cells and colonies were selected as described earlier (Li et al., 2020). The entire genome of ZIKV/mCherry and JEV/eGFP was verified by Sanger sequencing.

### Viruses and virus titrations

Infectious viruses were rescued from plasmid constructs by transfection into BHK-21J (pShuttle-YF17D, pShuttle-YFV/mCherry, pShuttle-JEV/eGFP and pShuttle-DENV2/mCherry) and Vero E6 (pShuttle-ZIKV/mCherry) cells using TransIT-LT1 Transfection Reagent (Mirusbio) following the manufacturer’s instructions (Li et al., 2020; Mishra et al., 2020). Upon onset of cytopathic effect (CPE), the recombinant viruses were subsequently passaged on their corresponding cell lines to generate virus stocks. DENV2/mCherry was generated in C6/36 as described earlier (Li et al., 2020). The harvested supernatants were centrifuged at 2100g for 8 min, aliquoted, and stored at −80°C. To determine the stability of mCherry insert in the YF17D backbone, the reporter virus was passaged up to passage 6 on BHK-21J. Virus stocks of passage 4 and 5 were used for all the experiments performed in this study.

YFV/mCherry, DENV2/mCherry and JEV/eGFP were titrated on BHK-21J cells; ZIKV/mCherry on Vero cells. Briefly, cells were seeded in 96-well black plates with transparent bottom (Greiner Bio-One) in assay medium and incubated overnight at 37°C. The next day, cells were inoculated with serial 3-fold dilutions of the reporter viruses and further incubated for 3 days. Hereafter, cells were fixed with 4% formaldehyde for 30 min at room temperature (RT), and washed once with Dulbecco’s phosphate-buffered saline (DPBS, Gibco). The plates were air-dried in the dark for 30 min prior to analysis. Scanning and counting of fluorescent spots were performed in a CTL Fluorospot reader using Fluorospot and Biospot modes (Software series 6 Ultimate, Cellular Technology Limited). Z’ values of each viral dilution were determined using the formula below:

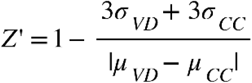

Here, σ and μ stand for standard deviation and mean of spots respectively, detected in either a certain virus dilution (VD) or cell control (CC).

### Viral RNA isolation and reverse transcription PCR (RT-PCR)

Total viral RNA was extracted from 0.1 ml virus stocks using Aurum Total RNA Mini Kit (Bio-Rad) and eluted in 50 μl elution buffer pre-heated to 70°C. The generation of cDNA and amplification of mCherry region was carried out using qScript XLT One-Step RT-PCR Kit (Quanta Bioscience). RT–PCR conditions were as follows: RT-step at 55°C for 20 min; initial denaturation at 94°C for 3 min; 40 cycles of amplification (denaturation at 94°C for 30 sec, annealing at 60°C for 45 sec, and elongation at 72°C for 50 sec), and final extension at 72°C for 10 min. The sequences of the primers were: YFV/mCherry-Forward (5′-GCAAATCGAGTTGCTAGGC-3′) and YFV/mCherry-Reverse (5′-CTTGAACACCTCTTGAAGG-3′). Each primer was used at a final concentration of 600 nM.

### Serum neutralization test (SNT^FLUO^)

A detailed protocol is available as Extended Data File. Briefly, sera were serially 1:3 diluted in triplicate in round-bottom 96-well plates and co-incubated with YFV/mCherry for 1 hour at 37°C. The YFV/mCherry inoculum titer was pre-determined by a titration experiment to result in 1500-2000 spot forming units. After one hour, the ‘serum-virus’ complexes are added in triplicate to pre-seeded cells and incubated at 37°C for one hour. Additionally, each 96-well plate contained six wells with

YFV/mCherry infected cells and six wells with uninfected cells serving as positive and negative controls, respectively. Fixation and imaging were performed similarly as described above in virus titration method section. The titers of nAbs protecting 50% (SNT_50_) and 90% (SNT_90_) of the cells from YFV/mCherry infection are determined by fitting the serum neutralization dilution curve that is normalized to the virus infection (100%) and cell control (0%) using Genedata Screener version 17.0.4. The limit of quantification was defined by the lowest dilution used in the assay; the limit of detection was determined by calculating the average neutralization titer multiplied by three times the standard deviation of 253 negative serum samples.

### Plaque reduction neutralization test

The PRNT assay was performed as described previously (Mishra et al., 2020). Briefly, BHK-21J cells were seeded in 12-well plates (Falcon) in seeding medium and cultured overnight at 37°C. The next day, triplicate serial dilutions of serum were co-incubated with YFV17D for 1h at 37°C prior to addition to BHK-21J cells. After 1h incubation, cells were washed twice with assay medium and overlaid with 2x Temin’s Modified Eagle Medium (Gibco) supplemented with 4% FBS and 0.75% sodium bicarbonate containing 0.5% low melting agarose (Invitrogen). The overlay was allowed to solidify at RT, cells were then cultured for 5 days at 37°C, fixed with 8% formaldehyde and visualized by staining with methylene blue. Plaques were manually counted and PRNT_50_ values were calculated by either serum dilution curve fitting using Genedata Screener version 17.0.4 or a nonlinear fitting algorithm (log(agonist) vs. response - Variable slope (four parameters)) in GraphPad version 8. The limit of quantification was defined by the lowest dilution used in the assay.

### Plaque reduction neutralization test (ISO 15189)

The certified PRNT assay was performed in 96-well plates (Falcon) using two-fold serum dilutions (ranging from 1:10 to 1:320) in six replicates. Briefly, serially diluted sera were co-incubated with pre-defined YFV17D titer for 1h at 37°C. Next, PS cells were added in DMEM supplemented with 5% FBS, 1% penicillin-streptomycin and 1% L-glutamine to the serum-virus mix and incubated during 3-4h. Cells were overlayed with 1.2% Avicel in DMEM supplemented with 5% FBS and incubated for 4 days at 37°C and 7% CO_2_, fixed with 3.7% formaldehyde and visualized by staining with Naphtol blue/black. Plaques were manually counted and PRNT_50_ values were calculated using the Reed-Muench method (Reed & Muench, 1938).

### Cell-based cytopathic (CPE) assay and CPE-based virus neutralization test (NT^CPE^)

The NT^CPE^ assay was performed as described previously (Mishra et al., 2020). Briefly, BHK-21J cells were plated in 96-well transparent plates (Corning) in seeding medium and incubated overnight at 37°C. Next day, triplicate serial dilutions of serum were co-incubated with 100 TCID_50_ of YF17D for one hour at 37°C and then added to the cells. After 5 days incubation at 37°C, each well was first visually inspected for the signs of CPE and then stained using MTS/PMS (Merck) for one to two hours at 37°C followed by absorbance reading at the 498 nm wavelength using a microtiter plate reader (Safire, Tecan). TCID_50_/ml was determined by serum dilution curve fitting using Genedata Screener version 17.0.4. The limit of quantification was defined by the lowest dilution used in the assay.

### Rapid fluorescent focus inhibition test (RFFIT, ISO 17025)

The RFFIT assay was performed as described previously (Roelandt et al, 2016). Briefly, sera, including positive and negative controls, were three-fold diluted (ranging from 1:9 to 1:243) in DMEM (Gibco) supplemented with 10% heat-inactivated FCS (Gibco). YFV strain French Neurotropic (National Collection of Pathogenic Viruses) was added at a dose of approximately 1.2 log TCID50 to the wells containing the diluted sera. Following this, BHK-21 cells were added to each well and incubated for 24h at 37°C and 5% CO2. Plates were fixed with 100% methanol at 4°C for 30 min. Infected BHK-21 cells were detected by an indirect immunofluorescence staining, using a primary mouse monoclonal antibody against the YFV envelope protein (Abcam) and a secondary Alexa488 fluor-conjugated goat anti-mouse IgG antibody (Molecular Probes). The number of foci with infected cells was counted under the fluorescence microscope. The SN titer was defined as the dilution of test serum that neutralized 50% of the virus (DIL50), calculated according to the Reed & Muench method (Reed and Muench, 1938).

### Ethics statement

Mouse and hamster sera used were sourced from historical samples from immunization experiments conducted at the KU Leuven Rega Institute, in accordance with institutional guidelines approved by the Ethical Committee of the KU Leuven, Belgium. Monkey sera were sampled from purpose-bred rhesus macaques (*Macaca mulatta*) housed at the Biomedical Primate Research Centre (BPRC, Rijswijk, The Netherlands) and vaccinated with either Stamaril® (Sanofi-Pasteur), or mock, upon positive advice by the independent ethics committee (DEC-BPRC), under project license DEC753C issued by the Central Committee for Animal Experiments, according to Dutch law. Human sera before and after vaccination with the YFV Vaccine Stamaril^®^ (Sanofi) were derived from a YF-17D vaccination study, approved by the ethical committee of the Medical Faculty, LMU Munich (IRB number 86-16).

### Serum samples and controls

Mouse, hamster, pig, monkey and human vaccinated for YF17D, JEV, ZIKV or DENV2 were collected for monitoring of other studies independent of this study. Monkey sera used for cross-validation of SNT^FLUO^ at the two Belgium reference centra at were collected from rhesus macaques vaccinated with licensed YF17D vaccine Stamaril® (Sanofi-Pasteur). Monkey anti-YF serum calibrated by a WHO International Reference Preparation (143 IU/ml) was used as a positive control (National & Control, 2011) both in PRNT and SNT^FLUO^. To inactivate complement, all sera were heat-inactivated by incubation at 56°C during 30 min prior to use.

### Statistical Analysis

GraphPad Prism (GraphPad Software, version 8) was used for all statistical evaluations. The number of animals or humans and number of replicate experiments that were performed is indicated in the figure legends. Correlation studies were performed using linear regression analysis with Pearson’s correlation coefficient and Bland-Altman analysis. Values were considered statistically significantly different at P values of ≤0.05.

## Data availability

The datasets generated and/or analyzed during the current study are available from the corresponding authors upon reasonable request.

## Acknowledgements

We thank Prof. Peter Bredenbeek (LUMC, The Netherland) for providing BHK-21J and Vero E6 cells and Prof. Andres Merits (University of Tartu, Estonia) for sharing cDNA of ZIKV strain. We also thank Javarappa Mahadesh Prasad Arkalagud for providing serum samples from ZIKV-vaccinated mice. Dr. Ernst Verschoor and Dr. Babs Verstrepen (Biomedical Primate Research Centre, BPRC Rijswijk, The Netherlands) are acknowledged for providing non-human primate sera. This project has received funding from the Research Foundation Flanders (FWO) under the Excellence of Science (EOS) program (No. 30981113; VirEOS), the European Union’s Horizon 2020 research and innovation program (No. 733176, RabydVax), KU Leuven intramural funding (IOF Hefboom; HB/13/010 and C3; C32/16/039), the German research foundation (Grant No. 391217598 and SFB/TR-237-B14 Grant No. 369799452).

## Author contributions

designed experiments; T.V., K.D, H.J.T.

carried out experiments; M.R., J.P., C.C., VS, L.H., J.T., J.M., K.G.

analyzed data; M.R., J.M., T.V., H.J.T.

provided advice on the interpretation of data; K.D., K.K.A., S.V.G.

wrote the original draft with input from co-authors; M.R., H.J.T.

wrote the final draft; M.R., T.V., K.D., H.J.T

provided and facilitated access to essential samples; M.N., L.L-H., K.D.B., S.R., H.K., J.T-S.

supervised the study; K.D., H.J.T

acquired funding; L.C., J.N., K.D.

All authors approved the final manuscript

## Materials and Correspondence

Correspondence and requests for materials should be addressed to K.D. and H.J.T.

## Figures and Legends

**Figure S1.**
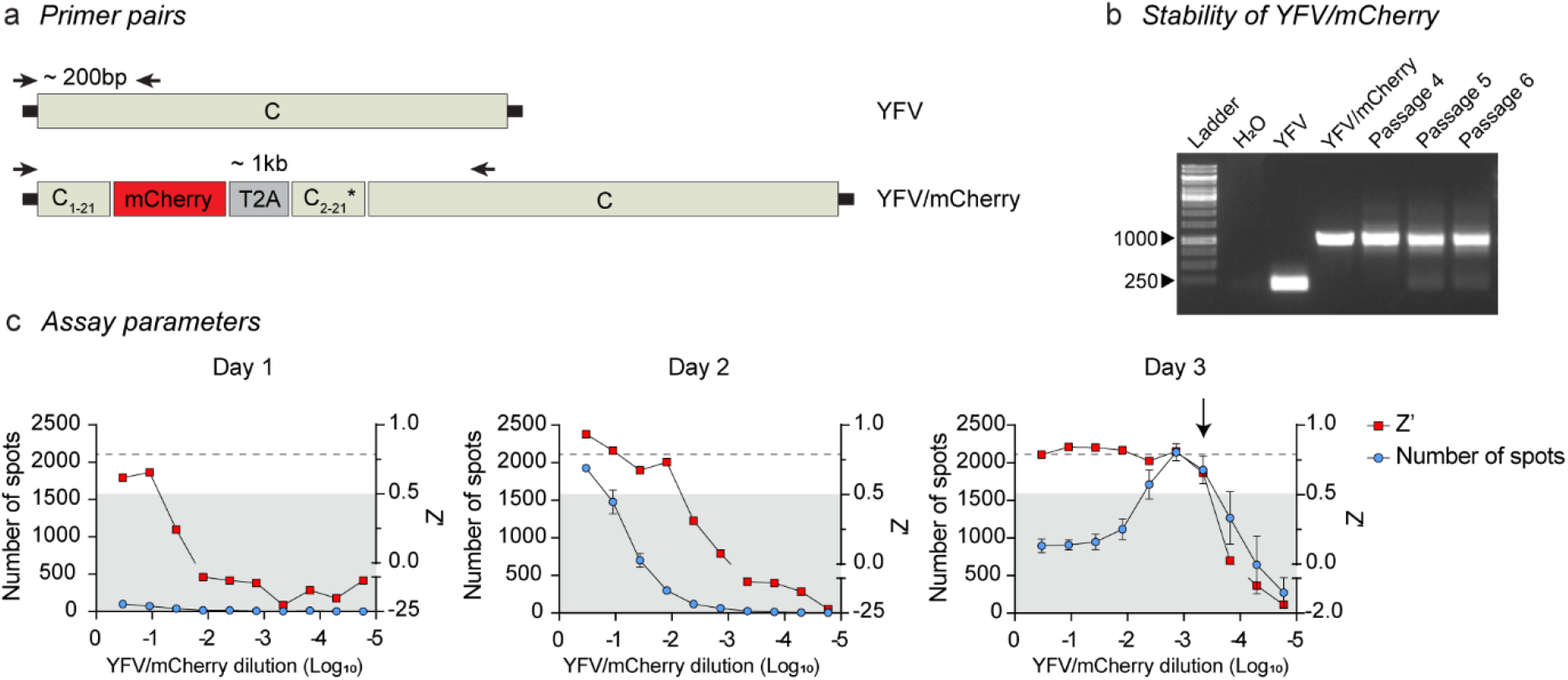
Optimization of SNT^FLUO^. **a**, Schematic representation of RT-PCR based detection of the mCherry insert. Arrows indicate primer binding sites on the viral genome. **b**, RT-PCR fingerprint of viral RNA extracted from infected BHK-21J supernatants of serially passaged YFV/mCherry (P4–P6). PCR amplicons of pShuttle-YFV and pShuttle-YFV/mCherry were amplified using the same primer pair and served as positive controls. H_2_O was included as a negative control. ladder, 1-kb DNA ladder. Data are from a single representative experiment. **c**, BHK-21J cells were infected with serially diluted YFV/mCherry. Spot counts (left Y-axis) and Z’ (right Y-axis) at indicated time points post infection. Grey area indicates Z’ values <0.5. Dashed line indicates saturation point of number of spots (∼2000 spots/well of a microtiter plate). Arrow in right panel indicates the highest Z’ value at the lowest virus dilution, immediately below a saturation point and was chosen for further assay development and validation. Data are the means ± standard deviations of three replicates.

**Figure S2.**
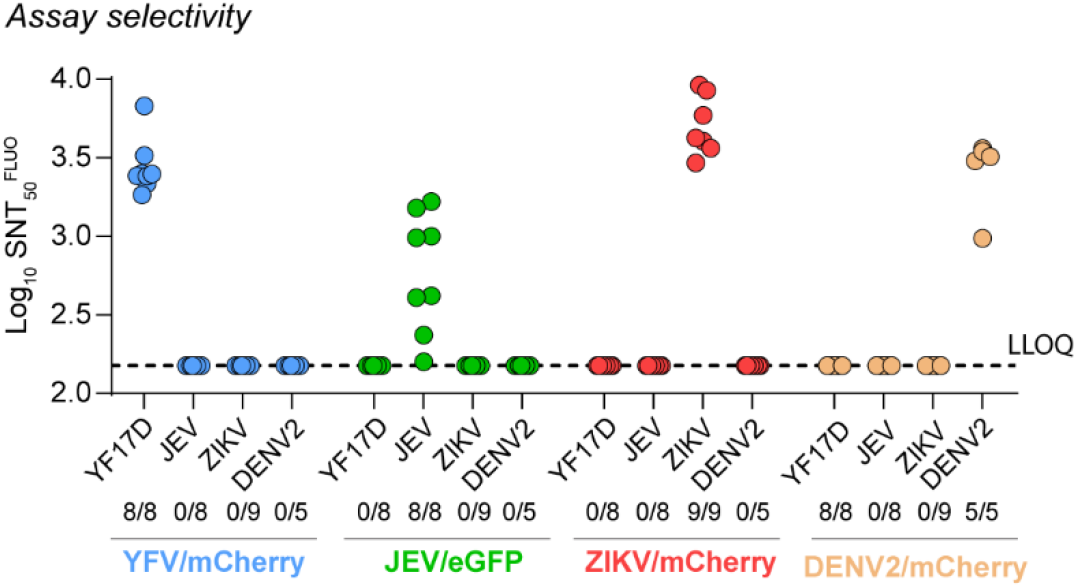
Selectivity of SNT^FLUO^ against other flaviviruses. Scatter plot of individual neutralizing titers (SNT_50_^FLUO^) for 30 serum samples from mice vaccinated with either YF17D (n=8), JEV (n=8), ZIKV (n=9) and DENV2 (n=5), and tested against YFV/mCherry, JEV/eGFP, ZIKV/mCherry and DENV2/mCherry. The number of seroconverted animals is shown as well. Data are the means of three replicates. LLOQ - Lower limits of quantification (dashed black line).

**Table S1.** SNT^FLUO^ validation by reference laboratories. Neutralizing titers of 1:10 serially diluted sera (1:10^1^ – 1:10^4^) from four non-vaccinated and six YF17D-vaccinated NHP. Data are the means of six (ITM and our laboratory) or two (Sciensano) replicates.

| Monkey ID | Day post-vaccination | log10 NT50-FLUO | log10 PRNT50 | log10 RFFIT50 |
| --- | --- | --- | --- | --- |
| R08086 | 42 | 4,2 | >3,1 | >2,9 |
| R08086 1:10 | 42 | 3,0* | 3,0* | 2,2* |
| R08086 1:100 | 42 | 1,6* | 1,9* | <1,8 |
| R08086 1:1000 | 42 | <1 | 1,7 | <1,8 |
| R08086 1:10000 | 42 | <1 | <1,6 | <1,8 |
| R09108 | 42 | >4,6 | >3,1 | >2,9 |
| R09108 1:10 | 42 | 3,3* | 3,0* | 2,7* |
| R09108 1:100 | 42 | 2,0* | 2,1* | 1,8* |
| R09108 1:1000 | 42 | <1 | 1,8 | <1,8 |
| R09108 1:10000 | 42 | <1 | <1,6 | <1,8 |
| R09131 | 42 | >4,6 | >3,1 | >2,9 |
| R09131 1:10 | 42 | 4,0* | 3,0* | 2,7* |
| R09131 1:100 | 42 | 2,4* | 2,4* | 1,8* |
| R09131 1:1000 | 42 | <1 | 1,8 | <1,8 |
| R09131 1:10000 | 42 | <1 | <1,6 | <1,8 |
| R04068 | 72 | 3,8 | >3,1 | >2,9 |
| R04068 1:10 | 72 | 2,8* | 3,0* | 2,4* |
| R04068 1:100 | 72 | 1,6* | 2,1* | <1,8 |
| R04068 1:1000 | 72 | <1 | 1,7 | <1,8 |
| R04068 1:10000 | 72 | <1 | <1,6 | <1,8 |
| R06081 | 72 | 4,5 | >3,1 | >2,9 |
| R06081 1:10 | 72 | 3,5* | >3,1 | 2,4* |
| R06081 1:100 | 72 | 2,0* | 2,4* | <1,8 |
| R06081 1:1000 | 72 | <1 | 1,7 | <1,8 |
| R06081 1:10000 | 72 | <1 | <1,6 | <1,8 |
| R12037 | 72 | 4,2 | >3,1 | >2,9 |
| R12037 1:10 | 72 | 3,3* | 2,8* | 2,6* |
| R12037 1:100 | 72 | 1,8* | 2,1* | <1,8 |
| R12037 1:1000 | 72 | <1 | <1,6 | <1,8 |
| R12037 1:10000 | 72 | <1 | <1,6 | <1,8 |
| R02006 | 42 | <1 | <1,6 | <1,8 |
| R07066 | 72 | <1 | <1,6 | <1,8 |
| R11050 | 72 | <1 | <1,6 | <1,8 |
| Pooled negative serum | 72 | <1 | <1,6 | <1,8 |

*\*used for correlation analysis*

